# Serial dependence in duration perception reveals reliability-weighted updating of the prior

**DOI:** 10.64898/2026.06.01.729229

**Authors:** Taku Otsuka, Hedderik van Rijn, Wouter Kruijne, Joost de Jong

## Abstract

Bayesian theories of perception propose that perceptual estimates result from the integration of prior beliefs (“priors”) with sensory input (“likelihood”), weighted by their reliability. While Bayesian theories assume that priors are continuously updated over time, empirical evidence for such reliability-weighted updating modulating sequential percepts remains lacking. Here, we leverage a behavioral phenomenon called serial dependence—in which perceptual judgments are attracted toward previous stimuli—to test a central prediction of Bayesian theories: that perceptual estimates are sequentially updated according to the reliability of successive stimuli. We used a duration reproduction task in which the reliability of perceived duration was manipulated via signal-to-noise ratio by embedding stimuli in dynamic white noise. Consistent with the prediction of Bayesian theories, serial dependence in perceived duration was enhanced by increased reliability of previous stimuli and attenuated by increased reliability of current stimuli. Computational modeling revealed that changes in sensory noise (i.e., the width of the likelihood) can account for the reliability-dependent modulation of serial dependence. These findings provide empirical evidence for reliability-weighted updating, supporting a central prediction of Bayesian theories that prior information and sensory input are iteratively integrated to calibrate perceptual estimates.

## Introduction

Bayesian theories of perception formalize perceptual inference as the integration of prior beliefs (“priors”) and sensory input (“likelihood”) to form posterior estimates. This integration is assumed to be reliability-weighted, with more reliable (i.e., precise) information exerting greater influence on perceptual estimates. Extensive empirical work supports this principle in the integration of multiple *concurren*t sensory signals (Alais & Burr, 2004; Ernst & Banks, 2002; Jacobs, 1999; Knill & Saunders, 2003) and of sensory input with fixed priors (Cicchini et al., 2012; Duhamel et al., 2023; Jazayeri & Shadlen, 2010; Körding et al., 2004; Maaß et al., 2021; Ueda et al., 2022). Another assumption of Bayesian theories is that perception is continuously updated over time, such that recent perceptual history serves as a prior (as the Bayesian dictum goes, “today’s posterior becomes tomorrow’s prior”) (de Jong et al., 2021; Petzschner et al., 2015). Behaviorally, this is reflected in *serial dependence*, in which perception of a current stimulus is attracted toward the immediately preceding stimulus (Cicchini et al., 2014, 2024; Fischer & Whitney, 2014; Manassi et al., 2023; Pascucci et al., 2023).

However, it remains unclear whether the reliability-weighted principles observed in the integration of concurrent input and of sensory input with fixed priors also apply to the perceptual updating across successive moments in time. From a Bayesian perspective, serial dependence reflects a form of reliability-weighted updating in which previous stimuli serve as a prior for current perceptual estimates. Under this account, the magnitude of attractive effects should depend on the relative reliability of current and previous stimuli: reliable previous stimuli increase the influence of prior information, whereas reliable current stimuli decrease it. Serial dependence therefore provides a critical test of a key prediction of Bayesian theories: that perceptual estimates are sequentially updated according to the reliability of successive stimuli.

Despite its theoretical importance, findings are mixed as to whether serial dependence is modulated by the reliability of both current and previous stimuli. Several studies have reported reliability-dependent modulation in serial dependence that is consistent with predictions of Bayesian models (Bergen & Jehee, 2019; Cicchini et al., 2018). However, these studies did not systematically examine the joint influence of current and previous reliability. By contrast, studies that explicitly tested this joint influence have yielded results that deviate from the predictions: serial dependence was primarily modulated by current reliability, with little or no influence from previous reliability (Ceylan et al., 2021; Gallagher & Benton, 2022).

One possible reason for these mixed findings is that previous studies have mostly focused on low-level visual features such as orientation (Bergen & Jehee, 2019; Ceylan et al., 2021; Cicchini et al., 2018; Fritsche et al., 2017, 2020; Gallagher & Benton, 2022; Gekas & Mamassian, 2025; Pascucci et al., 2019). In these domains, attractive serial dependence often coexists with repulsive serial effects associated with low-level sensory adaptation (Fischer & Whitney, 2014; Fritsche et al., 2017, 2020; Gekas et al., 2019; Manassi et al., 2018; Pascucci et al., 2019). Critically, manipulations intended to change stimulus reliability can also affect adaptation-induced repulsive effects (Gekas & Mamassian, 2025; Manassi et al., 2018; Pascucci et al., 2019), thereby confounding the interpretation of reliability-dependent changes in attractive serial dependence.

Duration perception, by contrast, provides a clearer test case for evaluating reliability-weighted updating across percepts. Serial dependence in perceived duration is well documented (Bertolasi et al., 2025; Chen et al., 2023; Cheng et al., 2024; de Jong et al., 2021; Dyjas et al., 2012; Sierra et al., 2022; Taatgen & van Rijn, 2011; Togoli et al., 2021; Wang et al., 2023), and attractive effects typically outweigh repulsive effects at short timescales relevant for serial dependence (e.g., the immediately preceding trial; Hayashi et al., 2015; Hayashi & Ivry, 2020; Heron et al., 2012, 2013; Maarseveen et al., 2017, 2018, 2019), reducing potential confounds from repulsive effects when manipulating stimulus reliability. Moreover, duration perception has been successfully modeled within Bayesian computational frameworks (Acerbi et al., 2012; Cicchini et al., 2012; de Jong et al., 2021; Glasauer & Shi, 2022; Jazayeri & Shadlen, 2010; Maaß et al., 2021; Petzschner et al., 2015; Roach et al., 2017; Sadibolova & Terhune, 2022; Shi et al., 2013), providing a formal basis for testing reliability-weighted principles.

Here, we capitalized on serial dependence in duration perception to test a central prediction of Bayesian theories: that perceptual estimates integrate current and previous information according to their reliability. Participants performed a duration reproduction task in which the reliability of perceived duration was manipulated by varying the signal-to-noise ratio (SNR) of a stimulus embedded in dynamic white noise. From a computational perspective, sensory noise (i.e., the width of the likelihood), which partially determines the relative weighting of prior and sensory information, can be most intuitively modulated by changing stimulus SNR. However, SNR manipulations may also influence the representation of duration itself (i.e., the mean of the likelihood), through modulations of stimulus contrast (Karsilar et al., 2026; Matthews et al., 2011), potentially producing reliability-dependent biases. To dissociate between these possibilities, we fitted and compared computational models within a Kalman filter framework that provides a canonical implementation of iterative Bayesian updating.

## Results

Participants performed a duration reproduction task in which they reproduced the duration of the presentation of a Gabor stimulus (Figure 1A). Stimulus duration had four levels (600, 750, 900, or 1050 ms). To manipulate the reliability of perceived duration (i.e., the width of sensory likelihood reflecting the duration), we embedded the Gabor stimulus in dynamic visual white noise (Figure 1B), creating two levels (high vs low reliability). Reliability was manipulated by varying the signal-to-noise ratio (SNR), either by changing Gabor contrast or noise opacity.

**Figure 1.**
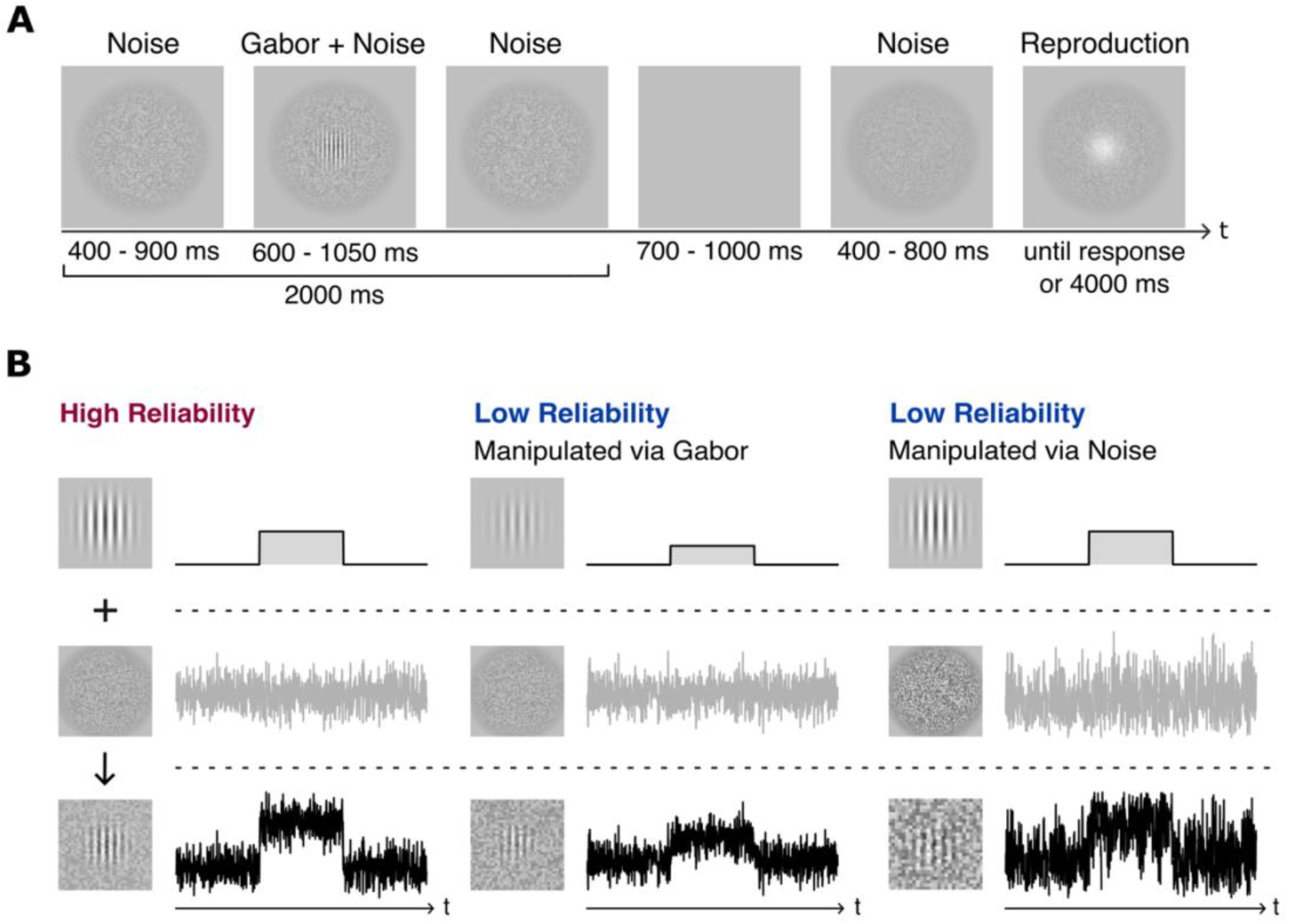
Schematics of the trial sequence and reliability manipulation. (A) Participants reproduced the duration of a Gabor stimulus embedded in dynamic white noise. (B) Stimulus reliability was manipulated by changing the signal-to-noise ratio (SNR), either by varying Gabor contrast or noise opacity, yielding high- and low-reliability stimuli. Stimuli do not reflect the exact stimulus parameters and are shown for illustrative purposes only.

### Reduced stimulus reliability increases response variability and reliance on prior expectations

Before analyzing serial dependence, we first assessed whether the two SNR manipulation methods (Gabor contrast vs. noise opacity) produced comparable behavioral effects. Across analyses, the manipulation methods showed no significant main effects or interactions (all *ps* > .31), suggesting that both methods affected behavior in a similar manner. We therefore aggregated data across the two manipulation methods in subsequent analyses of serial dependence.

A well-established prediction of Bayesian theories is that reduced sensory reliability—corresponding to a broader sensory likelihood—leads to an increased reliance on prior expectations. In duration perception, this should manifest as a stronger regression-to-the-mean effect for reduced-reliability stimuli, whereby perceived durations are pulled toward the mean of the earlier perceived durations, resulting in a flatter slope between reproduced and stimulus durations. Thus, we quantified this effect using the regression slope and found that it was flatter for low-reliability stimuli than for high-reliability stimuli (β = 44.74, t(18085.47) = 16.19, *p* < .001), indicating stronger regression to the mean (Figure 2A). Importantly, the regression slope in the low-reliability condition was significantly greater than zero (β = 18.64, t(49.61) = 6.08, *p* < .001), indicating that participants could still perform the task with low-reliability stimuli. These results are consistent with Bayesian predictions and indicate that our SNR manipulation successfully reduced sensory reliability, thereby increasing reliance on prior expectations.

**Figure 2.**
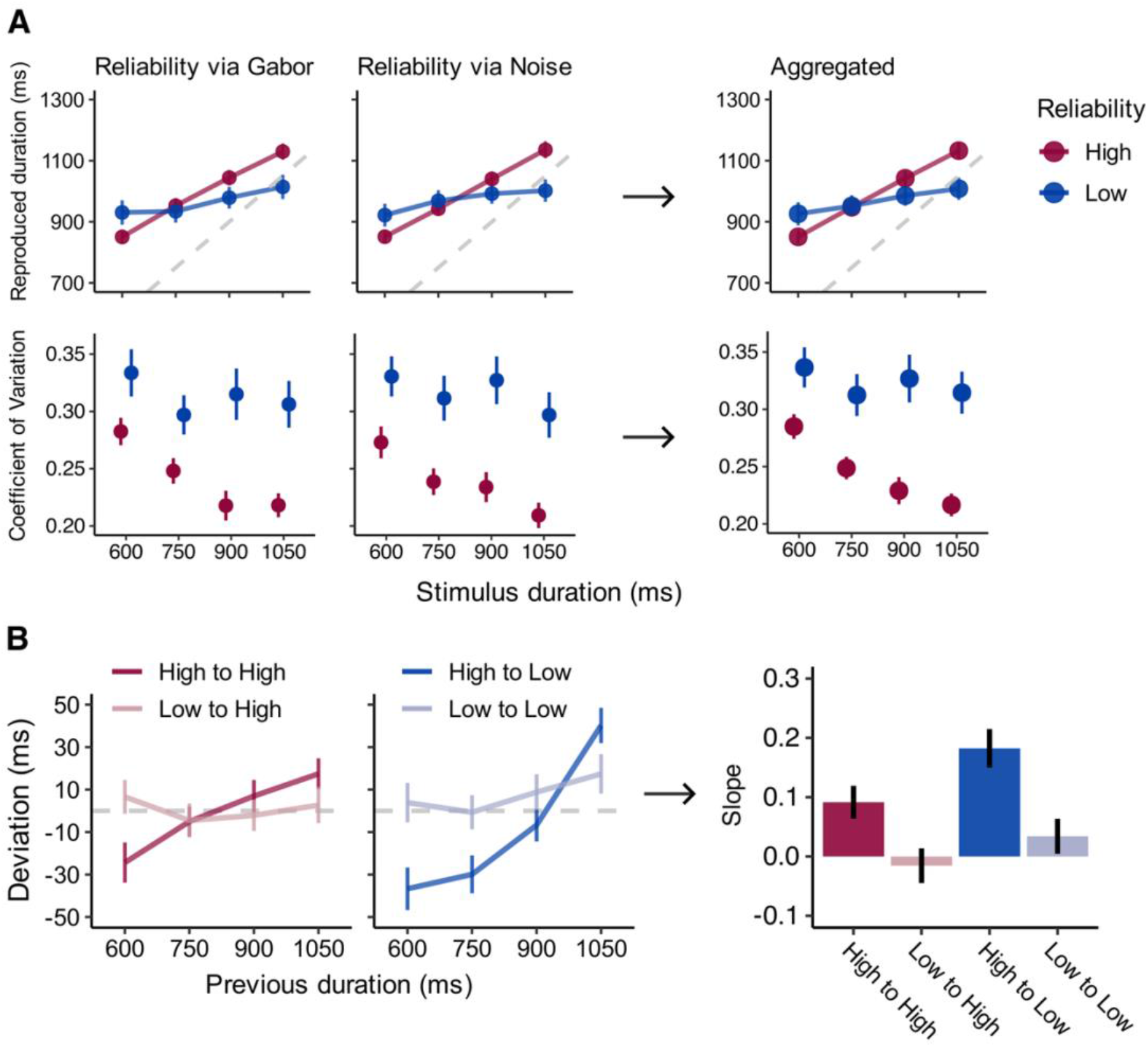
Behavioral effects of stimulus reliability. (A) Top panels show reproduced duration as a function of stimulus duration for each SNR manipulation method and for the aggregated data. Slopes were flatter for low-reliability stimuli, indicating a stronger regression-to-the-mean effect. Bottom panels show the coefficient of variation (CV), indicating greater response variability for low-reliability stimuli. Because no significant differences were observed between SNR manipulation methods, data were aggregated for subsequent analyses. (B) Serial dependence effects as a function of previous and current stimulus reliability. Left panels show trial-wise deviations plotted against previous (i.e., n-1) stimulus duration for each reliability condition. Right panel shows the corresponding regression slopes, indicating that attractive serial dependence increases with previous reliability and decreases with current reliability. Red and blue denote high and low current reliability, respectively; lighter shades indicate low previous reliability. Data show group means ± SEM.

Another expected consequence of reduced sensory reliability is that responses become more variable. We therefore quantified the response variability using the coefficient of variation (CV), which indexes variability relative to the mean response. As expected, CV was larger for low-reliability stimuli than for high-reliability stimuli (F(1,49) = 27.73, *p* < .001, η^2^_p_ = .36) (Figure 2A), providing converging evidence that our manipulation reduced sensory reliability. This increase was unlikely to reflect mean differences alone, as CV remained higher under low reliability at 750 ms, where mean responses were closely matched.

Together, these results confirm that our SNR manipulation modulated sensory reliability as intended, validating the key assumption of our experimental design.

### Current and previous reliability differentially enhance and attenuate the influence of prior information

A key prediction of Bayesian updating is that the influence of prior information depends on the reliability of both current and previous stimuli. Specifically, attractive serial dependence is expected to increase with previous (i.e., n-1) reliability and decrease with current reliability. Accordingly, we categorized trials by previous and current reliability into four conditions (High-to-High, Low-to-High, High-to-Low, and Low-to-Low). Serial dependence was quantified for each condition as the regression slope between the trial-wise deviations from participant-level means in reproduced duration and the stimulus duration on the previous trial (see Eqs. 1–2), with positive slopes indicating attractive effects toward the previous duration.

Consistent with the Bayesian predictions, serial dependence was modulated by the current and previous stimulus reliability (Figure 2B). Attractive effects were stronger when the previous stimulus was more reliable (F(1,49) = 12.76, *p* < .001, η^2^_p_ = .21) and weaker when the current stimulus was more reliable (F(1,49) = 4.97, *p* = .030, η^2^_p_ = .09). The interaction between previous and current reliability was not significant (F(1,49) = 0.67, *p* = .418, η^2^_p_ = 0.01), indicating that these effects were largely additive (see Figure S3 for a supplemental parameter profiling analysis of this interaction).

As predicted, attractive effects were strongest in the High-to-Low condition, and pairwise comparisons (Holm–Bonferroni corrected) revealed that this condition differed significantly from the Low-to-High (t(49) = 4.15, *p* < .001) and Low-to-Low conditions (t(49) = 3.39, *p* = .005). The other comparisons did not reach significance (High-to-Low vs. High-to-High: t(49) = 2.25, *p* = .080; Low-to-Low vs. Low-to-High: t(49) = 1.23, *p* = .443; Low-to-Low vs. High-to-High: t(49) = 1.20, *p* = .443; Low-to-High vs. High-to-High: t(49) = 2.45, *p* = .065). A complementary analysis with a linear mixed-effects model also showed that both current and previous reliability modulated serial dependence, as reflected in significant interactions with previous duration (current reliability: β = −6.86, t(17882) = −2.61, *p* = .009; previous reliability: β = 13.01, t(17882) = 4.96, *p* < .001). In summary, these results provide clear evidence for reliability-weighted updating across trials, whereby reliable previous inputs enhance the influence of prior information, whereas reliable current inputs attenuate it.

### A Bayesian updating model reveals sensory noise as the underlying mechanism of reliability-dependent attractive serial dependence

While the behavioral results support reliability-weighted updating, they do not specify which computational mechanism gives rise to these effects. To examine this, we used a Kalman filter model as a minimal implementation of Bayesian updating (Figure 3A; see Kalman filter model). Our primary hypothesis was that the reliability-dependent serial dependence would arise from changes in sensory uncertainty (i.e., sensory noise), which determine the relative weighting of prior and sensory information. However, because the reliability manipulations could also shift the representation of duration itself (i.e., sensory bias), we considered this as an alternative account. Within the Kalman filter framework, these candidate mechanisms are formalized as distinct components of sensory processing: sensory uncertainty is captured by the sensory noise parameter (*σ*_*m*_), corresponding to the variance of the likelihood, whereas shifts in encoded duration are captured by the sensory bias parameter (𝛥), corresponding to an offset in the mean of the likelihood.

**Figure 3.**
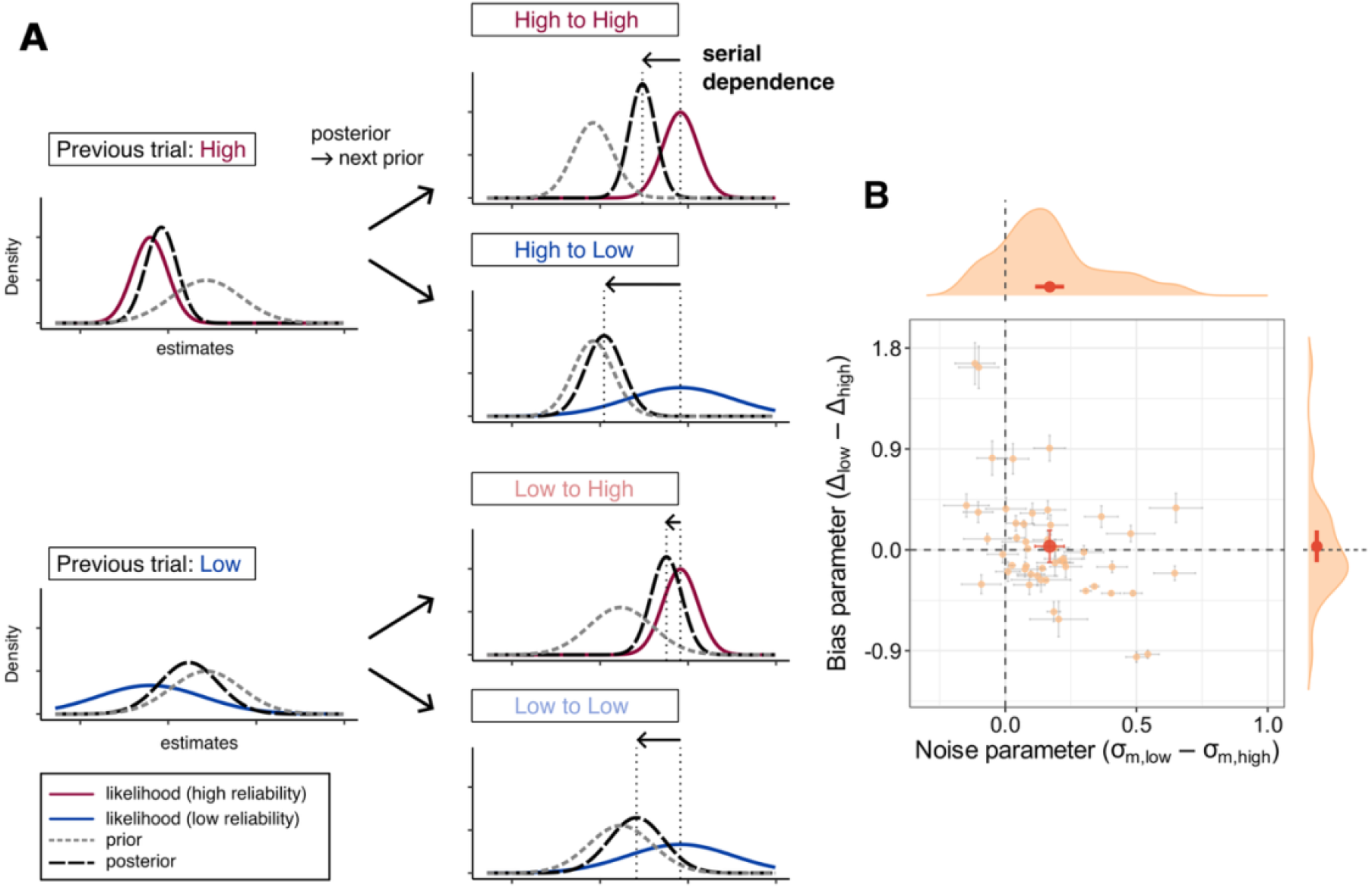
A Bayesian updating model and parameter estimates of sensory noise and sensory bias. (A) Schematic illustration of sequential Bayesian updating implemented by a Kalman filter. On each trial, the posterior estimate is carried over as the prior for the next trial and combined with the current sensory likelihood to form a new posterior. The four panels show how different combinations of previous and current reliability (high vs. low) affect the relative weighting of prior and likelihood, giving rise to reliability-dependent serial dependence. This schematic is for illustrative purposes and assumes that only sensory noise (i.e., the width of the likelihood) varies with reliability. (B) Differences in parameters of the full model between low- and high-reliability conditions. Differences in sensory noise (*σ*_*m*,low_ − *σ*_*m*,high_) and sensory bias (Δ_low_ − Δ_high_) are shown for each participant. Each point represents the mean parameter estimate across repeated fitting runs, and error bars indicate 95% confidence intervals across runs. Density plots show the distribution of parameter differences across participants. Red points and error bars indicate the group means and their 95% confidence intervals across participants. The noise parameter was significantly higher for low-reliability stimuli, whereas the bias parameter showed no systematic differences.

To assess whether the overall reliability effects are reflected in changes in sensory noise or sensory bias, we first fitted a full model in which both parameters were allowed to vary between high- and low-reliability stimuli. Model parameters (sensory noise: *σ*_*m*,high_, *σ*_*m*,low_; sensory bias: Δ_high_, Δ_low_) were estimated for each participant and compared at the group level. We found that the reliability effects were primarily captured by changes in sensory noise rather than sensory bias (Figure 3B). Specifically, sensory noise was higher for low-reliability stimuli than for high-reliability stimuli (*t*(49) = 6.14, *p* < .001, Cohen’s *dz* = 0.87), whereas sensory bias did not significantly differ between reliability conditions (*t*(49) = 0.45, *p* = .655, Cohen’s *dz* = 0.06), indicating that reduced stimulus reliability increases sensory uncertainty without inducing systematic shifts in sensory estimates.

To further determine which computational properties of the full model account for the observed reliability-dependent modulation in serial dependence, we compared three reduced models (see Reduced models): a “null model”, in which stimulus reliability did not affect sensory processing; a “noise model”, in which reliability selectively affected sensory noise; and a “bias model”, in which reliability selectively affected sensory bias. For each model and participant, model parameters were estimated by fitting reproduced durations, and two complementary Bayesian Information Criteria (BIC) were subsequently computed to evaluate model performance: BIC based on reproduced durations, which reflects both systematic and idiosyncratic components of behavior, and BIC_dev_ based on trial-wise deviations (as defined in Eq. 1), which isolates serial dependence effects from idiosyncratic reproduction biases. When model performance was evaluated using BIC based on reproduced durations, the bias model was most frequently selected (50% of participants, 25/50), followed by the noise model (36%, 18/50) and the null model (14%, 7/50). In contrast, when performance was evaluated using BIC_dev_, the noise model was most frequently selected (50%, 25/50), compared with the bias model (26%, 13/50) and the null model (24%, 12/50) (see Figure S2 for model comparison results). This pattern indicates that sensory noise accounts for the systematic reliability effects in serial dependence, whereas sensory bias primarily captures idiosyncratic effects of SNR on an individual’s reproduction behavior.

Altogether, the results of the full and reduced models point to a dissociation between sensory noise and sensory bias: sensory noise accounts for reliability-dependent attractive serial dependencies, whereas sensory bias primarily reflects idiosyncratic reproduction tendencies. Qualitative inspection of model predictions further supported this conclusion (Figure 4). Critically, the noise model qualitatively reproduced the key behavioral patterns—including stronger regression to the mean, increased response variability, and reliability-dependent modulation in serial dependence—whereas the null and bias models failed to capture these patterns. These findings indicate that reliability-dependent serial dependence is better explained by changes in sensory noise than by systematic shifts in encoded duration.

**Figure 4.**
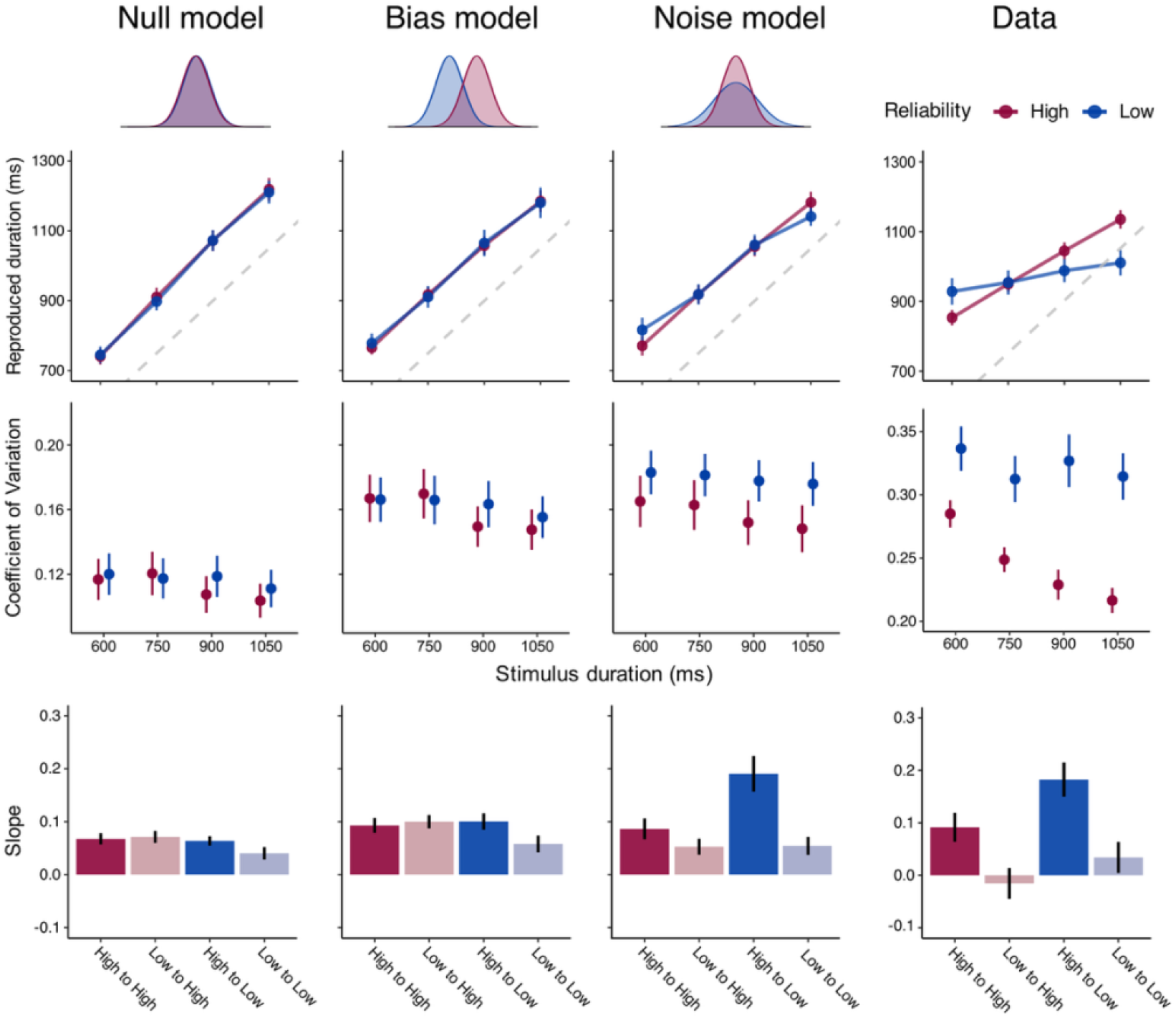
Model predictions and comparison with behavioral data. Model predictions for the null, bias, and noise models, alongside behavioral data (replotted from Figure 2). Top panels show reproduced duration as a function of stimulus duration, middle panels show the coefficient of variation (CV), and bottom panels show serial dependence effects quantified as regression slopes. Small schematics above each model illustrate the corresponding sensory likelihood. The noise model qualitatively reproduces the key behavioral patterns, whereas the null and bias models fail to capture them. Plotting conventions are the same as in Figure 2.

## Discussion

The present study assessed serial dependence in duration perception—a sequential effect in which perceived duration is biased toward previous stimuli—to test a central but unconfirmed prediction of Bayesian theories: that perceptual systems update their estimates across successive percepts in a reliability-weighted manner. By manipulating the reliability of successive stimuli, we show that attractive serial dependence is enhanced by increased previous reliability but attenuated by increased current reliability, consistent with Bayesian predictions. Computational modeling indicates that these effects can be accounted for by systematic changes in sensory noise (i.e., the width of the likelihood). Together, these findings demonstrate reliability-weighted updating, in which prior information is iteratively integrated with sensory input to calibrate perceptual estimates.

A key finding of the present study is that the influence of previous stimuli depends not only on the reliability of the *current* input but also on that of the *previous* input. Bayesian theories predict that the influence of prior information should depend on the uncertainty of prior and sensory input, with prior uncertainty inherited from the reliability of the previous input. While studies on serial dependence have consistently reported the influence of current reliability, evidence for the influence of previous reliability is lacking. For example, Cicchini et al. (2018) suggested that serial dependence is modulated by stimulus reliability in line with an ideal Bayesian observer model, and van Bergen & Jehee (2019) showed that fMRI-decoded sensory uncertainty predicts serial dependence in a Bayesian manner. However, these studies did not examine all reliability transitions: Cicchini et al. (2018) tested only conditions in which previous and current reliability were matched, whereas van Bergen & Jehee (2019) focused on cases in which they differed, leaving it unclear whether the observed effects truly reflect reliability-weighted updating or are driven solely by current reliability. To address this issue, Ceylan et al. (2021) and Gallagher & Benton (2022) manipulated all four reliability transitions as in the present study. They found an effect of current reliability but no effect of previous reliability, contrary to the Bayesian account.

The discrepancy between the present and previous findings may reflect methodological challenges in isolating reliability-dependent modulation in serial dependence. Prior work has focused on orientation perception, in which reliability is manipulated through stimulus features such as contrast (Pascucci et al., 2019), spatial frequency (Ceylan et al., 2021; Cicchini et al., 2018), or orientation bandwidth (Gallagher & Benton, 2022). While effective in altering reliability, these manipulations also affect early sensory processing, including tuning properties (Aghajari et al., 2020; Ayzenshtat et al., 2016; Broderick et al., 2022) and sensory adaptation (Crowder et al., 2006; Weigelt et al., 2011). Sensory adaptation poses an interpretational difficulty as it can produce repulsive serial effects that bias perception *away from* previous stimuli, potentially confounding the reliability-dependent changes in attractive serial dependence (Gekas & Mamassian, 2025). Critically, attractive and repulsive effects can coexist and operate over similar timescales in orientation perception (Fischer & Whitney, 2014; Fritsche et al., 2017, 2020; Gekas et al., 2019; Manassi et al., 2018; Pascucci et al., 2019): while attractive effects are typically strongest for the immediately preceding stimulus (Fritsche et al., 2020; Gekas et al., 2019), repulsive effects can also arise from a single brief exposure to a stimulus (e.g., 50 ms; Can & Collins, 2025) and extend across several to tens of trials (Fritsche et al., 2020; Gekas et al., 2019; Gekas & Mamassian, 2025). These findings indicate that the two effects may not be easily disentangled in analyses, such that apparent reliability-dependent changes in attractive serial dependence may partly reflect—or be masked by—changes in adaptation-related repulsive effects.

The present study was designed to reduce these potential confounds from early sensory processing, including adaptation-related repulsive effects, by leveraging duration perception. While low-level sensory parameters can affect duration processing (Bruno et al., 2011, 2015; Bruno & Johnston, 2010; Eagleman & Pariyadath, 2009; Johnston et al., 2006; Karsilar et al., 2026; Kruijne et al., 2021), duration perception is generally associated with relatively later stages of the processing hierarchy, such as frontoparietal networks (Harvey et al., 2020; Mondok & Wiener, 2023; Nani et al., 2019; Protopapa et al., 2019). Despite strong evidence that duration adaptation occurs after prolonged exposure to repeated durations (Heron et al., 2012; Li et al., 2017; Maarseveen et al., 2017), even these effects are linked to higher-level areas such as the parietal cortex (Hayashi et al., 2015, 2018; Hayashi & Ivry, 2020; Li et al., 2023). Crucially, attractive and repulsive effects are more temporally dissociable than in orientation perception: attractive effects are strongest for the immediately preceding trial and decay rapidly over the next few trials (e.g., within ∼1–3 trials; de Jong et al., 2021; Roseboom, 2019; Taatgen & van Rijn, 2011; Wang et al., 2023; Zimmermann & Cicchini, 2020), whereas repulsive effects typically emerge after repeated exposure to the same duration, often requiring tens of repetitions (Hayashi & Ivry, 2020; Heron et al., 2012; Maarseveen et al., 2019), with single exposures being insufficient to induce reliable adaptation effects (Li et al., 2017). These properties of duration perception allow a clearer assessment of reliability-dependent modulation in serial dependence by reducing sensory confounds from low-level visual processes and minimizing repulsive effects at the timescale relevant for attractive serial dependence.

Building on these advantages, we manipulated the reliability of the perceived duration by varying signal-to-noise ratio (SNR), which modulated sensory uncertainty: reduced SNR increased both regression to the mean and response variability, consistent with Bayesian predictions that increased uncertainty enhances reliance on prior information (Cicchini et al., 2012). Moreover, the two different SNR manipulations (Gabor vs. noise) produced comparable results, indicating that the observed effects cannot be attributed to differences in absolute stimulus intensity or Gabor contrast.

To further examine the computational mechanism underlying the observed reliability effects, we conducted model-based analyses using a Kalman filter framework (Petzschner et al., 2015). The results indicate that changes in sensory noise (i.e., the width of the sensory likelihood) provide a parsimonious account of the observed reliability effects. This is consistent with Bayesian accounts in which stimulus reliability is reflected in the precision of likelihood and thus determines the relative weighting of prior and sensory information. In contrast, sensory bias (i.e., the mean of the likelihood) appears to capture idiosyncratic tendencies in reproduction. Although sensory bias may reflect the influence of stimulus contrast or intensity on perceived duration (Bruno & Johnston, 2010; Karsilar et al., 2026; Kruijne et al., 2021; Matthews et al., 2011), such effects, if present, are not consistent across individuals and unlikely to account for the systematic reliability effects observed here. The Kalman filter framework may also help interpret the absence of a significant interaction between previous and current reliability in modulating serial dependence. In principle, such an interaction can be predicted by the model, but its predicted magnitude depends on the process noise parameter *q*. In Figure S3, we present a parameter profiling analysis illustrating this dependency. At the level of process noise estimated in our sample, the Kalman filter model predicts a much attenuated interaction effect, which could explain the lack of a significant interaction in the behavioral data. Future studies with larger samples, more reliability conditions, or experimental manipulations of process noise may provide a more sensitive test of the predicted interaction.

While the Kalman filter model provides a general and parsimonious framework for sequential Bayesian updating (Daw et al., 2006; Petzschner et al., 2015; Wolpert et al., 1995), it does not capture the full complexity of the underlying neurocognitive mechanisms and has several limitations. First, it cannot account for repulsive serial effects, which may coexist with attractive serial dependence. Second, it assumes that all past information is summarized in a single prior, effectively collapsing longer-term influences into the immediately preceding estimate, even though longer-term and short-term effects may be partially dissociable. More elaborate models are therefore required for a detailed mechanistic account. For example, the ACT-R-based memory model (Taatgen & van Rijn, 2011), the two-state Bayesian model (Glasauer & Shi, 2022), or the mixed-Gaussian updating model (Wang et al., 2023) provide finer descriptions of contextual effects in duration perception. In orientation perception, several two-stage models have been proposed in which attractive and repulsive effects arise from distinct mechanisms (Fritsche et al., 2020; Gekas & Mamassian, 2025; Pascucci et al., 2019). How such models can account for reliability-dependent effects remains an important question for future research.

The present study shows that information from previous trials is integrated with current input; however, the study remains agnostic about the processing stage at which this integration occurs. Serial dependence has been attributed to multiple stages, including perceptual stages (Cicchini et al., 2021; Fischer & Whitney, 2014), working memory (Fritsche et al., 2017), decisional stages (Akaishi et al., 2014; Pascucci et al., 2019), and motor responses (Wu et al., 2026). Moreover, signals arising at higher-level stages can propagate back to influence early sensory processing (Cicchini et al., 2021), suggesting that these stages interact recurrently rather than operate independently. Although the present data cannot determine their relative contributions, several observations help constrain the interpretation. First, attractive serial dependence in duration perception has been observed in tasks that do not involve reproduction (de Jong et al., 2021; Zimmermann & Cicchini, 2020), suggesting a locus before motor stages. Second, our Kalman filter framework assumes that stimulus reliability affects sensory noise at the encoding stage and that the prior for the next trial is inherited directly from the posterior, independent of the actual response, making motor responses unlikely to be the primary locus of the reliability-weighted integration observed here. Determining how these different processing stages contribute will require further research.

Taken together, the present results provide empirical evidence for a central prediction of Bayesian theories: reliability-weighted updating of the prior over time. Building on previous findings that recent perceptual history is incorporated as a dynamically updated prior (Cicchini et al., 2014, 2018; de Jong et al., 2021), we show that this updating depends on the reliability of both previous and current inputs. These findings extend the scope of reliability-weighted integration beyond its established role in the integration of multiple concurrent signals and sensory input with fixed priors, demonstrating that perception is iteratively updated through reliability-weighted integration to calibrate perceptual inference over time.

## Methods

### Participants

Fifty-one first-year students at the University of Groningen participated in exchange for partial course credit (38 females; aged 18–25 years). All participants reported normal or corrected-to-normal vision, normal hearing, and no history of psychiatric or neurological conditions. On the basis of criteria set by the Ethics Committee of the Faculty of Behavioural and Social Sciences at the University of Groningen, the study (PSY-2425-S-0097) was exempt from full review. The study was conducted in accordance with the NETHICS code, and written informed consent was obtained before participation. One participant was excluded from further analysis based on a predefined performance criterion.

### Apparatus

Stimulus generation and presentation were controlled by OpenSesame 4.0 (Mathôt et al., 2012) with the PsychoPy backend (Peirce et al., 2019). Visual stimuli were displayed on a 27-in. Iiyama ProLite G2773HS monitor (1920 × 1080 resolution, 100 Hz). Participants were seated approximately 60 cm from the display in a dimly lit room.

### Stimuli

Visual stimuli were presented on a mid-gray background (50% intensity). Each stimulus consisted of a circular noise patch (10.1° diameter) with raised-cosine edges. The patch contained dynamic uniform noise, which was updated on every frame. Noise contrast was fixed at 70%. A Gabor patch (2.85° diameter, 3.4 cycles/deg) was presented at the center of the noise patch. Stimulus reliability (signal-to-noise ratio; SNR) was manipulated using two SNR manipulation methods: either via the Gabor contrast or the noise opacity (Figure 1B). In the Gabor-manipulation condition, SNR was varied by changing the contrast of the Gabor stimulus (25% for low reliability and 100% for high reliability) while noise parameters were fixed (70% contrast and 50% opacity). In the noise-manipulation condition, Gabor parameters were fixed (100% contrast and 100% opacity), and SNR was manipulated by changing the opacity of the noise patch (80% for low reliability and 50% for high reliability) while noise contrast remained constant (70%). All intensity values are expressed as percentages of the maximum PsychoPy rendering parameters.

### Procedure

Participants performed a duration reproduction task (Figure 1A). Each trial began with an inter-trial interval (ITI) during which a gray background was presented for 500–800 ms (uniform distribution). The stimulus interval then began with the presentation of the noise patch, which remained on the screen for 2000 ms. A Gabor stimulus was presented within the noise patch for one of four durations (600, 750, 900, or 1050 ms). The onset of the Gabor stimulus occurred 400–900 ms after the onset of the noise patch (uniform distribution), and the noise patch remained visible after the Gabor offset until the end of the 2000-ms interval. Following the stimulus interval, a blank inter-stimulus interval (ISI) of 700–1000 ms (uniform distribution) was presented. The noise patch was then presented again, followed by a white Gaussian patch that served as the reproduction cue. Participants reproduced the perceived duration of the Gabor stimulus by pressing the space key to terminate the white patch after they felt the same amount of time had elapsed since its onset. The reproduction display remained until a response was made or 4000 ms had elapsed, after which the next trial began automatically.

Stimulus duration had four levels (600, 750, 900, and 1050 ms), and stimulus reliability had two levels (high and low) (Figure 1B). Stimulus reliability (i.e., SNR) was manipulated either through Gabor contrast or noise opacity, which were presented in separate blocks. The experiment consisted of six blocks, with three blocks for each SNR manipulation method. Gabor-manipulation and noise-manipulation blocks alternated, and the order of the first block was counterbalanced across participants. Each block contained 64 trials, comprising all combinations of duration and reliability repeated eight times. Thus, participants completed 384 trials in total. For 21 participants tested earlier in the study, each block contained 56 trials (seven repetitions per condition), as the number of repetitions was increased during data collection. Trial order within each block was determined using a de Bruijn sequence to ensure balanced transitions between stimulus conditions. Before the main experiment, participants completed a short practice session using the same stimuli (approximately 12 trials).

### Analysis

Trials with reproduced durations shorter than the lower bound defined by the 1.5 × interquartile range (IQR) criterion were removed. The IQR was computed from the distribution of reproduced durations pooled across all trials and participants, and the lower bound was defined as Q1 − 1.5 × IQR. Trials with reproduced durations longer than 3900 ms were also excluded to remove timeout responses. In total, 0.3% of trials were excluded based on these criteria. To exclude participants who did not perform the duration reproduction task as instructed, we fitted a linear regression between stimulus duration and reproduced duration for each participant using data from high-reliability trials. Participants whose regression slope was negative were excluded from further analyses. This criterion resulted in the exclusion of 1 out of 51 participants.

### Manipulation checks

To verify that our SNR manipulation affected stimulus reliability, we analyzed reproduced durations and response variability. Reproduced durations were analyzed using a linear mixed-effects model implemented in R with the *lme4* package (Bates et al., 2015) and *lmerTest* package (Kuznetsova et al., 2017). Reproduced duration on each trial served as the dependent variable. Fixed effects included stimulus duration (mean-centered and divided by 100), stimulus reliability (low vs. high), SNR manipulation method (Gabor vs. noise), and all interactions. Stimulus reliability and SNR manipulation method were effect-coded (−0.5, 0.5 for low/Gabor and high/noise conditions, respectively). Random intercepts and random slopes of participant for stimulus duration were included. This analysis tested whether the regression-to-the-mean effect—quantified as the slope relating stimulus duration to reproduced duration—varied as a function of stimulus reliability and whether this effect depended on the SNR manipulation method.

Response variability was quantified using the coefficient of variation (CV), defined as the standard deviation of reproduced durations divided by their mean. CV was computed separately for each participant and condition defined by stimulus duration, stimulus reliability, and SNR manipulation method. The resulting CV values were analyzed using a repeated-measures ANOVA with reliability, SNR manipulation method, and stimulus duration as within-participant factors.

### Serial dependence

To quantify serial dependence effects independently of baseline reproduction biases, we analyzed trial-wise deviations from condition-specific mean responses. For each participant, mean reproduced durations were first computed separately for each combination of stimulus duration, stimulus reliability, and SNR manipulation method. The deviation on trial *n* was then defined as:

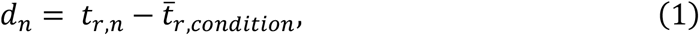

where *t*_*r,n*_ is the reproduced duration on trial *n*, and 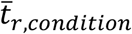 is the mean reproduced duration for the corresponding condition. By subtracting the condition-specific mean, this normalization removes systematic biases associated with the current stimulus (e.g., regression-to-the-mean effects), thereby isolating trial-by-trial fluctuations that can be attributed to sequential influences. Because the manipulation checks indicated no differences between the two SNR manipulation methods, data were collapsed across manipulation methods for subsequent analyses. Serial dependence was then quantified by regressing trial-wise deviations onto the stimulus duration of the preceding trial:

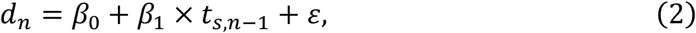

where *t*_*s,n*−1_ is the stimulus duration of the preceding trial. Regression slopes were estimated separately for each participant and for each combination of previous-trial and current-trial reliability (High to High, Low to High, High to Low, and Low to Low), using data from all stimulus durations. Positive slopes indicate attractive serial dependence toward the previous stimulus. The resulting slope estimates (*β*_1_) were analyzed using a repeated-measures ANOVA with previous and current stimulus reliability as within-subject factors. Pairwise comparisons were conducted using paired-samples t-tests, with p-values adjusted using the Holm–Bonferroni method.

In order to assess reliability-dependent modulation at the trial level, we also fitted a linear mixed-effects model in which trial-wise deviations were predicted by the stimulus duration of the previous trial and its interactions with current and previous stimulus reliability. Reliability factors were effect-coded (−0.5 for low, 0.5 for high), and the previous stimulus duration was centered (825 ms) and scaled by 100. A random intercept for participants was included.

### Modeling

#### Kalman filter model

To identify the computational mechanism underlying reliability-dependent modulation in serial dependence, we implemented a Kalman filter model that formalizes sequential Bayesian updating across trials. All computations were performed in log-duration space.

On each trial *n*, the observer’s belief about stimulus duration is represented by a Gaussian prior with mean *μ*_pri,*n*_ and variance 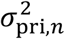 . The sensory measurement on trial *n*, denoted *x*_*m,n*_, is a noisy observation of the stimulus duration *t*_*s,n*_:

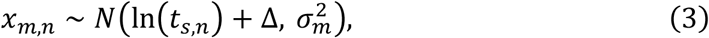

where the sensory bias parameter (Δ) captures a systematic shift in the mean of the sensory measurement, and the sensory noise parameter 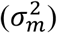 determines its variance.

Given this measurement, the prior is updated to a posterior distribution with mean *μ*_post,*n*_ and variance 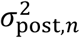 according to:

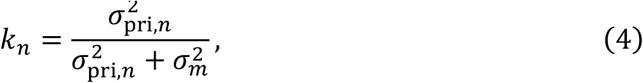

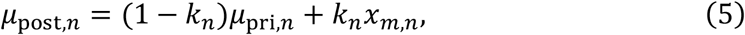

where *k*_*n*_ is the Kalman gain that determines the relative weighting of prior and sensory information, such that higher sensory noise reduces the weight assigned to current sensory input and increases reliance on the prior.

The prior for the next trial is inherited from the posterior on the current trial:

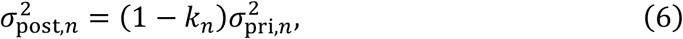

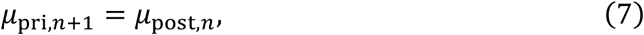

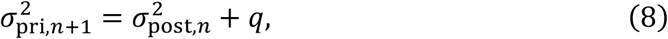

where *q* represents process noise that captures the increase in uncertainty across trials.

Finally, the reproduced duration is defined as the mean of the corresponding log-normal distribution derived from the posterior:

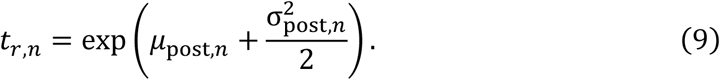

### Full model

Our behavioral results indicate that stimulus reliability modulates the extent to which prior information influences reproduced durations. Within the Kalman filter framework, such effects can arise from two distinct components of sensory processing. First, the sensory noise parameter 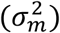 determines the uncertainty (i.e., the variance) of the sensory measurement (Eq. 3), thereby influencing the Kalman gain that controls the relative weighting of prior and sensory information (Eq. 4). Second, the sensory bias parameter (𝛥) determines the systematic shift (i.e., the mean) of the sensory measurement (Eq. 3).

To examine how stimulus reliability affects these components, we first fitted a full model in which both sensory noise and sensory bias parameters were allowed to vary across stimulus reliability. In this model, separate sensory noise parameters (*σ*_*m*,low_ and *σ*_*m*,high_) and sensory bias parameters (Δ_low_ and Δ_high_) were estimated for low- and high-reliability stimuli, while a single process noise parameter *q* was estimated across conditions. Parameter estimation was repeated using 100 different random seeds for each participant to ensure robust convergence of the optimization procedure. For each participant, parameter estimates were averaged across seeds. Reliability-dependent differences in sensory noise and sensory bias parameters were then assessed across participants using paired samples t-tests. Because this model was introduced to examine reliability-dependent differences in parameter estimates rather than to evaluate model structures, it was not included in the model comparison.

### Reduced models

To further clarify the role of sensory noise and sensory bias, we compared three reduced models that differed in how stimulus reliability affected sensory processing.

The “null model” assumed that stimulus reliability does not affect sensory processing. In this model, a single sensory noise parameter 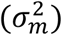 and a single sensory bias parameter (Δ) were estimated across all trials, serving as a baseline model.

The “noise model” assumed that reliability selectively modulates sensory uncertainty. Separate sensory noise parameters were estimated for low- and high-reliability stimuli (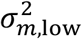 and 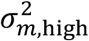), while the sensory bias parameter (Δ) was estimated across conditions. Because sensory noise determines the Kalman gain, this model implements reliability-weighted updating.

The “bias model” assumed that reliability selectively affects sensory bias. Separate bias parameters were estimated for low- and high-reliability stimuli (Δ_low_ and Δ_high_), while a single sensory noise parameter 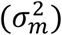 was estimated across conditions.

In all models, a single process noise parameter *q* was estimated.

### Model fitting

Model parameters were estimated separately for each participant by minimizing the residual sum of squares (RSS) between model predictions and observed data. For each candidate parameter set, the model simulated reproduced durations using the same stimulus sequence experienced by the participant. To capture the distribution of responses, we used a quantile-based fitting procedure. Trials were grouped according to the combination of current stimulus reliability, previous stimulus reliability, and previous stimulus duration, resulting in 16 condition cells. Within each cell, reproduced durations were summarized using the 10th, 30th, 50th, 70th, and 90th percentiles. The same summary statistics were computed for simulated data. Parameter optimization was performed using Particle Swarm Optimization, as implemented in the *pso* package (Bendtsen, 2022) in R. The search space was bounded as follows: sensory noise parameters ∈ [0.001, 2], sensory bias parameters ∈ [−2, 2], and process noise parameters ∈ [0.001, 2]. The initial values of the prior mean and variance were fixed across participants (see Figure S1 for overall goodness-of-fit across models).

### Model comparison

Model comparison was conducted to determine which computational mechanism best accounts for the observed reliability effects in serial dependence. For each participant and fitted model, responses were simulated using the estimated parameters and the same stimulus sequence experienced by the participant. Model performance was quantified using the RSS computed on quantile summaries, following the same procedure as in model fitting, and the Bayesian Information Criterion (BIC) was approximated from RSS assuming Gaussian residuals:

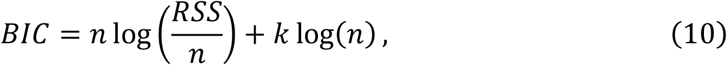

where *n* denotes the number of data points (*n* = 16 *cells* × 5 *quantiles* = 80), and *k* denotes the number of free parameters. The null model contained three parameters (*σ*_*m*_, Δ, *q*), whereas the noise model and the bias model each contained four parameters. We performed two complementary model comparisons that differed in the behavioral measure used to compute the quantile summaries and the resulting RSS. First, BIC was computed using reproduced durations. This measure captures overall behavioral patterns, including both systematic effects (e.g., regression to the mean and serial dependence) and idiosyncratic biases in reproduction. Second, an analogous deviation-based criterion (BIC_dev_) was computed using trial-wise deviations (Eq. 1), which remove condition-specific mean biases and isolate trial-by-trial fluctuations attributable to sequential dependencies. This measure therefore provides a more direct assessment of how well each model captures serial dependence. Notably, because model fitting was based on reproduced durations, BIC_dev_ should be interpreted as a penalized model-evaluation measure, rather than as a conventional maximum-likelihood-based BIC. Comparing model performance across these two measures allowed us to determine whether a given model accounts for general reproduction behavior or specifically for the reliability-dependent modulation in serial dependence.

## Data and code availability

Data and code are available in the Open Science Framework repository: https://osf.io/sk9qy

