## Supplementary Materials for "Serial dependence in duration perception reveals reliability-weighted updating of the prior"

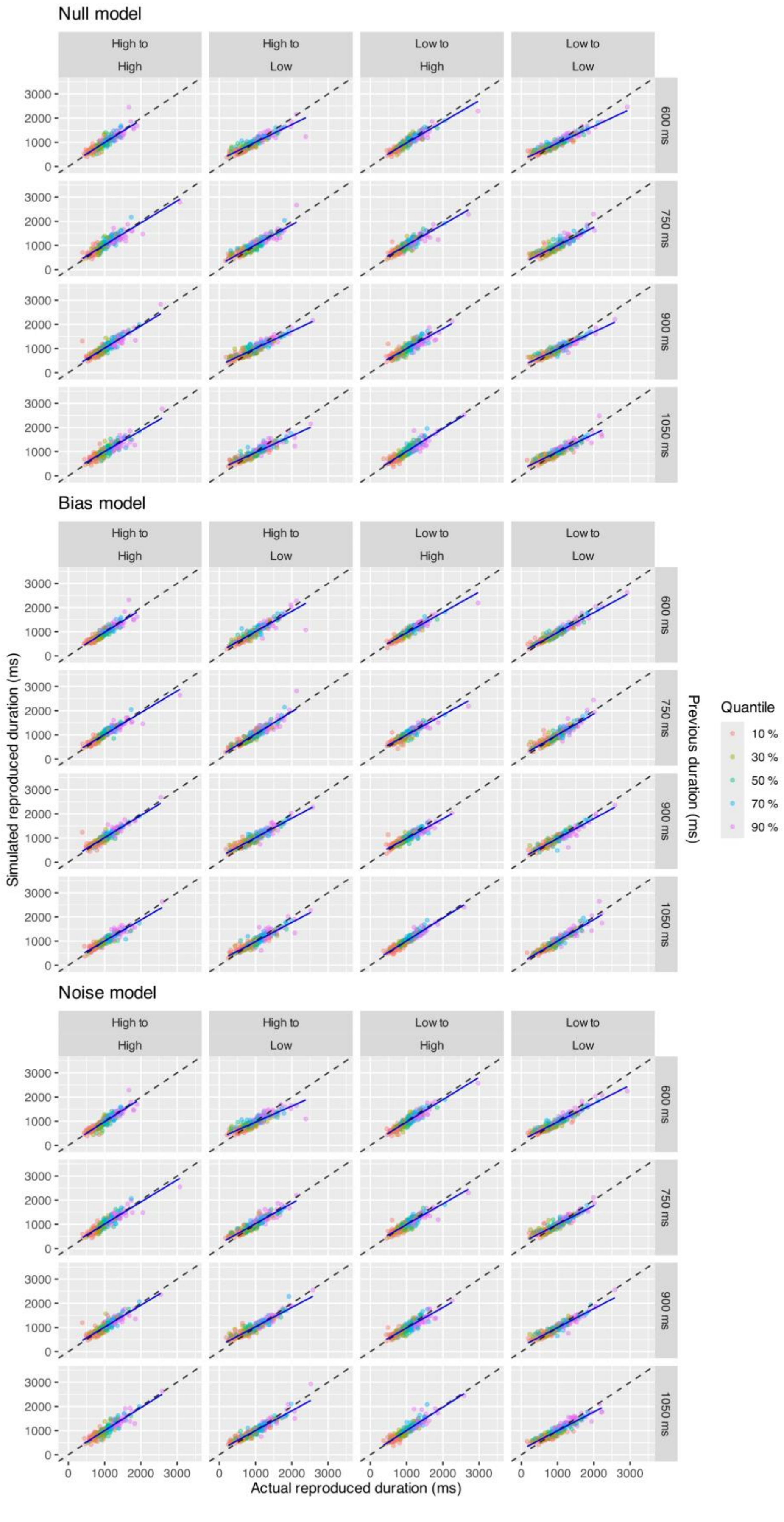

Null model
Bias model
Noise model
High to High
High to Low
Low to High
Low to Low
600 ms
750 ms
900 ms
1050 ms
0
1000
2000
3000
Simulated reproduced duration (ms)
Actual reproduced duration (ms)
Previous duration (ms)
Quantile
10 %
30 %
50 %
70 %
90 %

**Figure S1. Goodness-of-fit of the models.**

Scatter plots compare actual and simulated reproduced duration quantiles for three models (null, bias, and noise). Data are shown separately for each combination of previous reliability, current reliability, and previous stimulus duration. Colors indicate quantiles (10%, 30%, 50%, 70%, and 90%). Each point represents one participant's quantile estimate in a given condition. The dashed diagonal line indicates perfect agreement between actual and simulated values, and the blue line shows the linear fit.

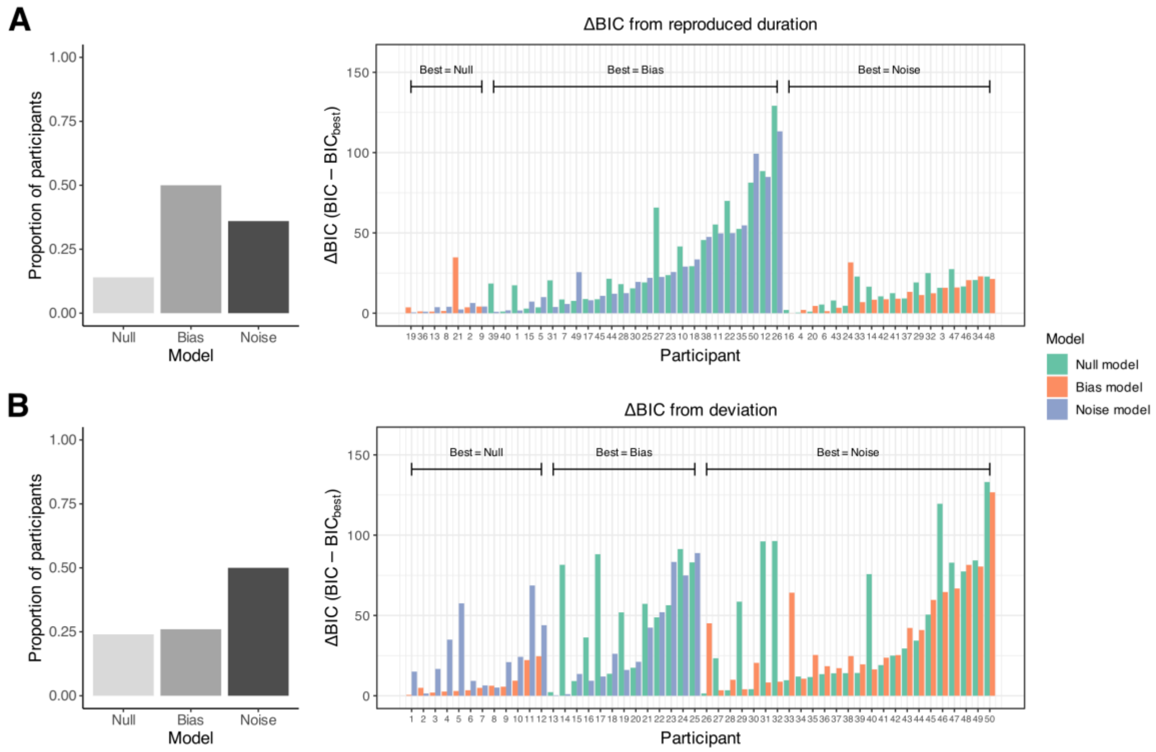


**Figure S2. Model comparison based on BIC and BIC$_{dev}$.**

Model performance was evaluated for three models (null, bias, and noise) using BIC computed from the residual sum of squares (RSS). (A) BIC computed from the RSS of reproduced duration quantiles. (B) BIC$_{dev}$, an analogous deviation-based criterion computed from the RSS of deviation quantiles. In each panel, the left subpanel shows the proportion of participants for whom each model was selected as the best (i.e., lowest BIC), and the right subpanel shows participant-wise ΔBIC values, defined as the difference from the best model within each participant. Larger values indicate poorer model performance relative to the best model. Participants are grouped by their best model (indicated by horizontal brackets) and, within each group, sorted by the difference between the best and second-best models. For consistency, participant indices on the x-axis correspond to the ordering derived from the deviation-based comparison in panel B and are shared across both panels.

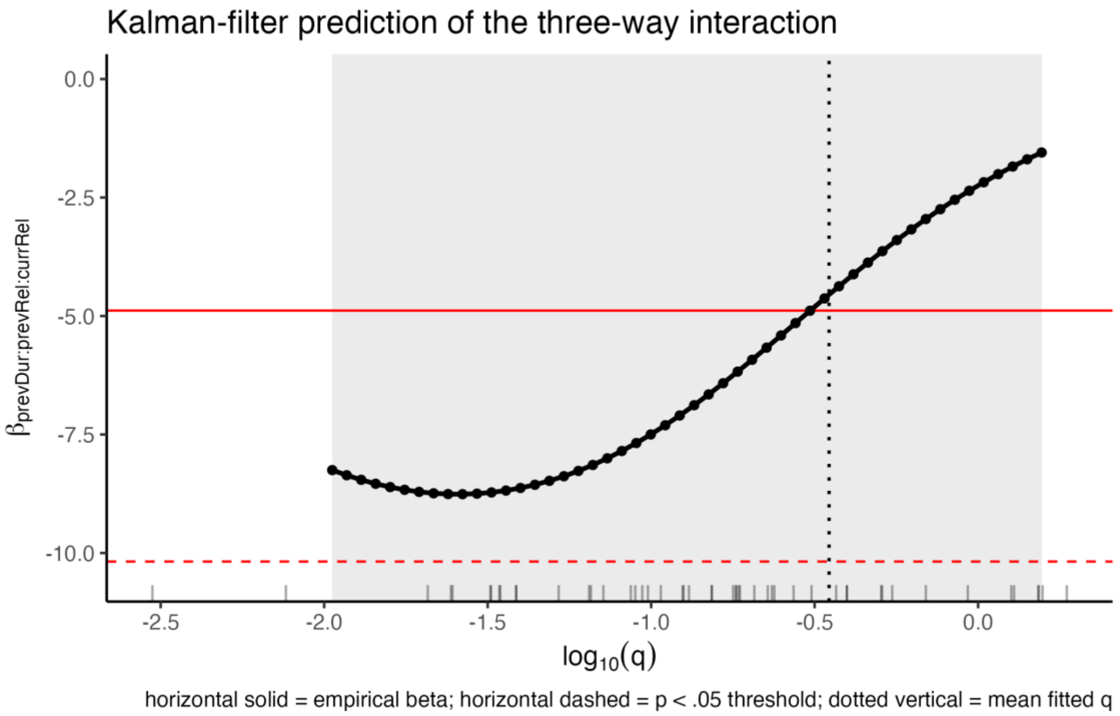


**Figure S3. Parameter profiling analysis of the Kalman filter prediction for the Previous Duration × Previous Reliability × Current Reliability interaction.**

To examine the Kalman filter model's prediction regarding the interaction effect between current and previous reliability on serial dependence, we conducted a parameter-profiling analysis focusing on the process noise parameter $q$. In this analysis, $q$ was varied across an empirically plausible range, defined as the 2.5th to 97.5th percentiles of the fitted $q$ estimates from the full model. For each value of $q$, model-predicted responses were generated using the actual stimulus sequences from the experiment, while retaining each participant's fitted sensory-noise and sensory-bias parameters from the full model. Trial-wise deviations were then computed in the same way as in the behavioral analysis. The fixed-effect $\beta$ for the Previous Duration × Previous Reliability

× Current Reliability interaction ($\beta_{prevDur:prevRel:currRel}$) was estimated using the same LMM structure as in the behavioral analysis: Deviation ~ prevDur * prevRel * currRel + (1 | subject).

The black line and dots show the model-predicted β as a function of $q$. The bottom rug shows participant-level fitted $q$. The dotted vertical line indicates the mean fitted $q$. The solid red horizontal line indicates the empirical beta, and the dashed red horizontal line indicates the critical $\beta$ magnitude required for the interaction to reach significance at $\alpha = .05$, computed as $|\beta_{\text{critical}}| = t_{.975,df} \times SE_{\text{empirical}}$, where $SE_{\text{empirical}}$ and the degrees of freedom were taken from the empirical LMM.

The profiling analysis shows that the magnitude of the interaction depends on $q$, with smaller $q$ values yielding more negative interactions. Notably, the empirical $\beta$ was close to the model-predicted $\beta$ around the mean fitted q, suggesting that the observed interaction is broadly consistent with the Kalman-filter prediction at the empirically estimated level of process noise. Moreover, across the empirically plausible range of $q$, the model-predicted $\beta$ remained below the critical $\beta$ magnitude required for significance. Thus, while the Kalman-filter model does predict an interaction between current and previous reliability, this interaction is attenuated at empirically plausible levels of process noise, suggesting that the nonsignificant interaction observed in the behavioral data does not contradict the Kalman-filter model.
